# PLZF limits enhancer activity during hematopoietic progenitor aging

**DOI:** 10.1101/504324

**Authors:** Mathilde Poplineau, Julien Vernerey, Nadine Platet, Lia N’guyen, Léonard Hérault, Michela Esposito, Andrew J. Saurin, Christel Guilouf, Atsushi Iwama, Estelle Duprez

## Abstract

PLZF (promyelocytic leukemia zinc finger) is a transcription factor acting as a global regulator of hematopoietic commitment. PLZF displays an epigenetic specificity by recruiting chromatin-modifying factors but little is known about its role in remodeling chromatin of cells committed towards a given specific hematopoietic lineage. In murine myeloid progenitors, we decipher a new role for PLZF in restraining active genes and enhancers by targeting acetylated lysine 27 of Histone H3 (H3K27ac). Functional analyses reveal that active enhancers bound by PLZF are involved in biological processes related to metabolism and associated with hematopoietic aging. Comparing the epigenome of young and old myeloid progenitors, we reveal that H3K27ac variation at active enhancers is a hallmark of hematopoietic aging. Taken together, these data suggest that PLZF, associated with active enhancers, appears to restrain their activity as an epigenetic gatekeeper of hematopoietic aging.

## INTRODUCTION

Transcription factors (TFs) play important roles during hematopoiesis, from stem cell maintenance to lineage commitment and differentiation (1, 2). It is now well admitted that sequence specific TFs in concert with epigenetic factors regulate gene expression during hematopoiesis (3, 4). For more than a decade, the progression of hematopoietic stem cells (HSCs) towards more differentiated cells has been thought to be accompanied by epigenetic reprograming (5, 6) and it appears that the structure of chromatin is essential for hematopoietic lineage specification (7, 8). However, the interdependent interactions between TFs and chromatin features during hematopoietic differentiation are still poorly understood. Studies in different systems have indicated that the chromatin of regulatory elements is in a pre-active state in stem cells and/or early progenitors before the transcriptional initiation, leading to the concept of “gene priming” (9). The priming is thought to be driven by a specific class of TFs called “pioneer TFs,” that are able to induce the early chromatin changes during the gene activation process (10).

PLZF, also known as Zbtb16, is a master transcriptional regulator with effects on growth, self-renewal and differentiation with a well-recognized role in hematopoietic, spermatogonial, mesenchymal and neural progenitor cells (11). Within the hematopoietic tissue, PLZF is involved in the production of numerous different hematopoietic and immune cells and regulates immune responses (12, 13). Its expression marks NKT-cell development (14) but also controls megakaryopoiesis (15) or invariant natural killer T cell effector functions (16). However, PLZF is not considered as a lineage-specific TF unlike lineage-instructive TFs such as PU.1 and C/EBPalpha (17–19). Indeed, PLZF inactivation in mouse models does not result in a specific blockage in the immature hematopoietic compartment (20), but induces subtle changes in HSC cell cycle progression that participate to the aging of the hematopoietic system (21). This aging-like phenotype in PLZF-mutant HSC was characterized by an increase in myeloid progenitor differentiation at the expense of the lymphoid progenitors and was correlated with the alteration of gene expression programs related to stem cell function and cell cycle (21).

PLZF has a recognized epigenetic function; it is probably through its chromatin activity that PLZF finely and precisely regulates transcriptional programs that mediate its biological functions. However, it remains unclear how PLZF causes changes in the existing epigenetic landscape (22). PLZF binds to multiple epigenetic cofactors that could be recruited and modify the chromatin landscape at the vicinity of PLZF chromatin localization (23–25). For instance, previous studies have shown that PLZF recruits HDAC complexes at targeted promoters and locally reduced histone acetylation (25, 26). In addition, we previously showed that presence of PLZF at chromatin is associated with the tri-methylation of lysine 27 of histone H3 (H3K27me3) enrichment at developmental genes (24, 27).

Here, we reveal a new chromatin function of PLZF in myeloid progenitors. By analyzing epigenomic landscape variations upon PLZF expression or inactivation, we discovered that PLZF inhibits acetylation of lysine 27 of histone H3 (H3K27ac) at enhancer regions that are already active. Moreover, the de-repression of the enhancer regions observed in PLZF-mutated granulocytic-monocytic progenitors (GMPs) is also observed in old GMPs. Thus, we propose that PLZF limits some aging features by restricting enhancer activity of genes involved in metabolic processes.

## MATERIAL AND METHODS

### Mice

The mouse model C57BL/6 Cd45-2 *Zbtb16^lu/lu^* was previously described in Vincent-Fabert et al. (21). For reconstitution 2 millions of total bone marrow were transplanted in lethally irradiated (8-8.5 Gy) C57BL/6 Cd45-1 mice. Reconstitution was monitored every 4 weeks and mice were sacrificed for GMP purification at 16 weeks post transplantation. Young (~2 months) and aged (~18 months) C57BL/6-Cd45.1 mice were purchased from Charles River Laboratories. B6-Cd45.1/Cd45.2 mice were bred and maintained in the CRCM mouse facility in accordance with our institutional guidelines for the use of laboratory animals and approved by the French authority (authorization number: MESR#5645).

### Purification of granulocyte monocyte progenitors (GMPs)

Tibias and femurs were crushed in PBS containing 3% of fetal calf serum (FCS). Red blood cells were lysed using ACK buffer (Gibco). Bone marrows were depleted in mature cells, expressing the lineage markers Cd5, B220, Cd11b, Gr-1, Ter-119 or 7-4, using the lineage cell depletion kit (Miltenyi Biotec). GMPs (Lineage^-^, Cd45^+^, C-Kit^+^, Sca-1^-^, Cd34^+^, FcγR^+^) were purified using the FACS Aria III cell sorter (Beckman Dickinson). Antibodies used for GMP staining are described in supplemental materials.

### Cell culture and lentiviral transduction

The 416b murine myeloid cell line expressing the surface marker Cd34 (a generous gift from B. Göttgens) was maintained at exponential growth in RMPI 1640, 10 % FCS and 1% penicillin/streptomycin. Cells were transduced with PLZF-FLAG-GFP or empty-FLAG-GFP lentiviral particles, 30 min at 2000 rpm, 32°C. After 48 h, GFP positive cells were purified on the FACS-ARIA II cell sorter (Beckman Dickinson).

### Luciferase Assay

416b cells were transfected using Amaxa^®^ Cell Line Nucleofactor^®^ kit C (Lonza), U937 program, with 2 μg of ***REN1LLA*** Luciferase control reporter and 4 μg of ***CD47*** enhancer-Firefly vectors. ***CD47 enhancer-FIREFLY*** vector was generated from the previously described construct ***CD47-E5*** (28). Luciferase activity was monitored 8 h after transfection using the Dual Luciferase Reporter Assay (Promega).

### Reverse-Transcription and PCR quantitative (RT-qPCR)

Total RNA was extracted using the RNeasy Plus Micro kit (Qiagen). cDNA was synthesized with the Transcriptor High Fidelity cDNA Synthesis Kit (Roche). PCR was done using Taqman probes (Mm00607939_s1 Actb and Mm01176868_m1 Zbtb16; Life Technologies) and Taqman Universal PCR Master Mix (Life Technologies) on the Fast 7500 Real-Time PCR system (Applied Biosystem). Relative expression levels were determined by the 2^^-ΔCT^ method using *Actb* as housekeeping gene.

### Chromatin immunoprecipitation (ChIP)

For histone mark ChIP-seq, purified GMPs or 416b cells were fixed with 1% of formaldehyde for 8 min. Reaction was quenched by adding 2 mM of glycine. ChIP procedures are developed in supplemental experimental procedure. ChIP-seq libraries were generated using the MicroPlex Library Preparation Kit (Diagenode) following the manufacturer’s instructions and analyzed on a 2100 Bioanalyzer system (Agilent) prior sequencing.

### Quantification of immunoprecipitated DNA

Quantification of immunoprecipated DNA was performed by quantitative real time PCR (qPCR). qPCR was performed using SsoAdvanced™ Universal SYBR^®^ Green Supermix (BIO-RAD) with the CFX PCR system (BIO-RAD). For enrichment quantification, Input Ct values were subtracted to ChIP Ct values and converted into bound value by 2^(-(Input Ct-ChIP Ct))^. Data are expressed as % of bound/input.

### Primer sequences are shown in supplemental materials

For normalization of ChIP-qPCR, Spike-in Drosophila Chromatin and Spike-in Antibody (Active Motif) were added to the ChIP reaction as a minor fraction of the IP reaction according to the manufacturer’s instructions. The immunoprecipitated Drosophila chromatin was quantified using Drosophila Positive Control Primer Set Pbgs (Active Motif), allowing the attribution of a spike-in factor to each sample, used for normalization.

### ATAC-seq

ATAC-seq was performed on 30 000 cells. Cells were washed twice with cold PBS and suspended in lysis buffer (10 mM Tris -HCl pH 7.4, 10 mM NaCl, 3 mM MgCl2, 0.1% IGEPAL CA-630). The same amount of cells was used for input preparation. Transposition and library preparation were done using the Nextera DNA Library Prep Kit (Illumina) according to the manufacturer’s instruction. Size selection was performed using the Blue Pippin^TM^ system.

### RNA-seq

Total RNA was extracted using the RNeasy Plus Mini kit (Qiagen). GMPs cDNA synthesis and library preparation were done by Beckman Counter Genomics using a TruSeq Stranded Total RNA with Ribo Zero Gold kit (Illumina). 416b cDNA was synthesized using a SMART-Seq v3 Ultra Low Input RNA kit for Sequencing (Clontech). ds-cDNA was fragmented using S220 Focused ultrasonicator (Covaris) and libraries were generated using a NEBNext Ultra DNA Library Prep kit (New England Biolabs).

### Sequencing and data processing

For GMPs, ChIP-sequencing was performed with a Next-seq500 sequencer (Illumina) using a 75-nt single-end protocol, at the Paoli Calmettes Institute Sequencing Facility (IPC, Marseille) and RNA-sequencing with a Next-seq500 sequencer (Illumina) using a 100-nt paired-end protocol, at Beckman Counter Genomics. For 416b cells, ChIP and RNA sequencing was performed with a HiSeq1500 sequencer (Illumina) using a 61-nt single-end protocol at Chiba University, Japan. Computational analyses are developed in supplemental experimental procedure. Profiles of histone ChIPseq and ATAC-seq signals were obtained by processing normalized bigwig files through the deepTools suite (v2.2.4)(29) (computeMatrix, plotProfile). For PLZF ChIP-seq, genomic distribution and annotation of PLZF peaks in 416b cell line were determined using HOMER software (v4.7.2; annotatePeaks module) (30). All sequencing data were visualized using the Integrative Genomics Viewer (IGV v2.3.92) (31). Biological functions associated with PLZF-ChIP peaks were determined using Genomic Regions Enrichment of Annotations Tool (GREAT v3.0.0) (32). The same tool was also used for assigning enhancers to the closest genes. The Gene Ontology (GO) biological processes associated with candidate genes were determined using g:Profiler (Reimand, NAR, 2016) tool with a p-value < 0.05 and by taking into account a background. For enhancers named “K27ac de novo”, “K27ac up”, “K27ac unchanged” (supplemental Figure S4) the background used was all active enhancer-associated genes contained in the 416b PLZF-Flag and 416b Empty vector conditions. For enhancers named “common” (Figure 5E), the background used was all active enhancer-associated genes contained in WT, *Zbtb16^lu/lu^*, Old and Young GMP conditions.

Biological functions associated with PLZF-ChIP peaks were determined using Genomic Regions Enrichment of Annotations Tool (GREAT v3.0.0) (32). The same tool was also used for assigning enhancers to the closest genes.

### Statistics

For two group analyses, we first check the normal distribution of each group (Shapiro-Wilk test) and then performed Welch’s t-test. Density plot profils’ p-values were obtained by performing paired Welch’s t-test on computeMatrix averaged signal matrices. Concerning Venn diagram analysis, overlap significances were computed by hypergeometric tests (phyper function, lower.tail=False). Concerning ChIP-qPCR and Luciferase assay, Mann Whitney test was used.

## RESULTS

### PLZF restricts active epigenetic marks in myeloid cells

To better understand PLZF activity in mouse myeloid cells, we profiled PLZF genomic occupancy in the murine myeloid 416b cell line (a cellular model for Cd34-positive progenitors). As PLZF-ChIPseq in murine cells was not successful, we generated a PLZF-expressing 416b cell line by transducing 416b cells with lentivirus containing PLZF-Flag and by selecting a stable expressing clone (**Figure 1A**). This stable cell line was first used to profile PLZF genomic occupancy using Flag antibody. In PLZF-Flag 416b cells, PLZF binds mostly intragenic regions (63.6%) with preference for intronic sequences and TSS regions, in accordance with previously reported genomic localization in human myeloid cells (27) (**Figure 1B**). Presence of PLZF on its previously described targeted genomic loci was confirmed by Integrative Genomics Viewer (IGV) visualization (**Supplemental Figure S1A**) and peak calling analysis. In line with our previous study (27), genes bound by PLZF-Flag in the myeloid progenitor cell line (**Table 1**) were enriched for Gene Ontology (GO) terms relating to cell cycle, DNA integrity and immunity. Remarkably, genes involved in protein acetylation-related GO terms were also enriched within the genes targeted by PLZF (**Figure 1C**), underlying a potential effect of PLZF on Histone lysine acetylation. To assess the epigenetic landscape of this cell line, we performed H3K27ac, H3K4me3, H3K4me1 and H3K27me3 ChIP-seq. Our analyses show that in PLZF-Flag 416b, PLZF was significantly associated with transcriptionally active histone marks. We found that more than 41% of PLZF peaks overlapped with H3K27ac and H3K4me1, 38.6% with H3K4me3. Overlap of PLZF peaks with the transcriptionally repressive histone mark H3K27me3 was low (2.9%) and not significant (**Figure 1D**).

**Figure 1:**
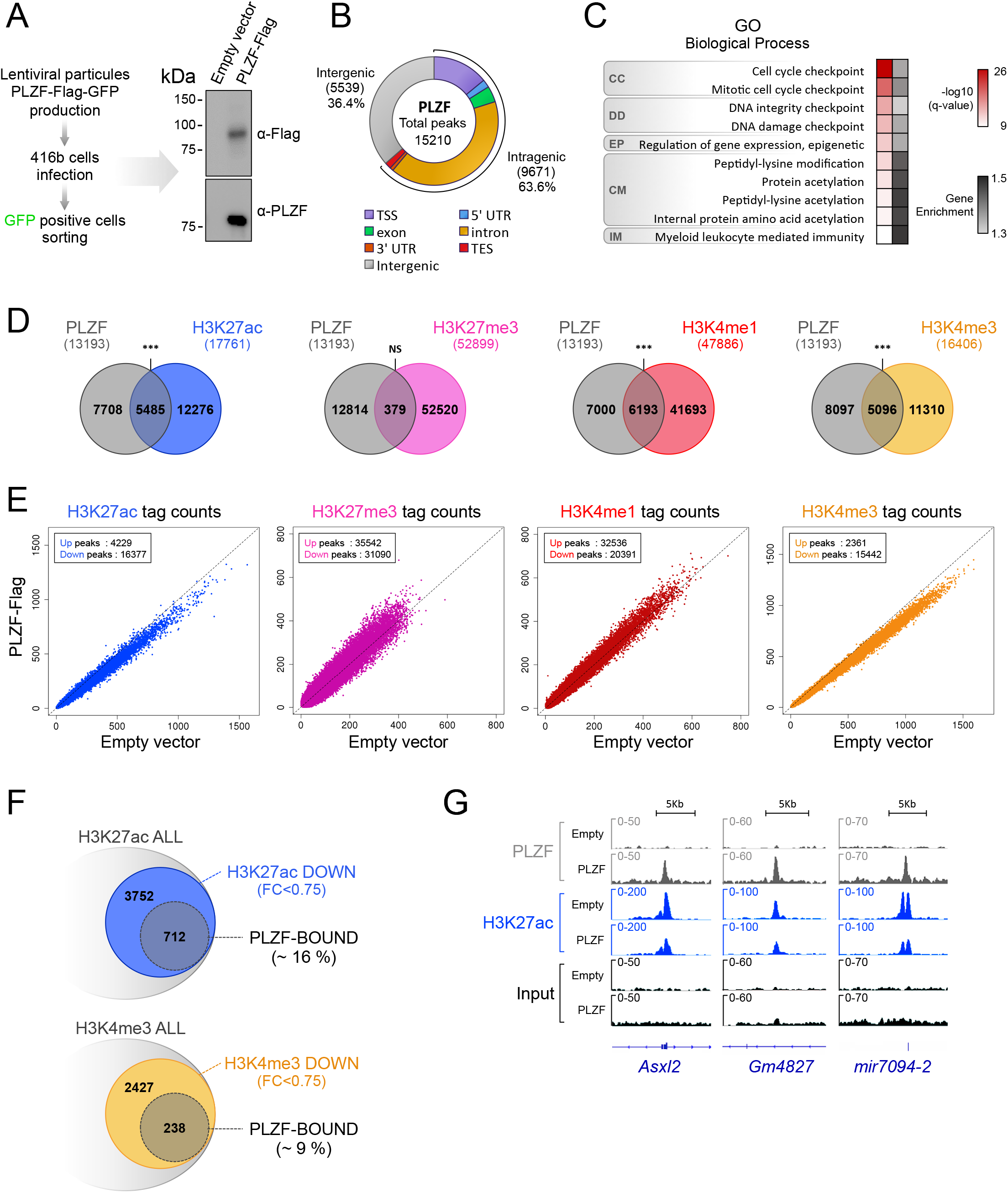
PLZF restricts active epigenetic marks in myeloid cells. (A) Experimental protocol design for engineering PLZF-Flag 416b cell line and immunoblot showing PLZF expression (using anti-Flag or anti-PLZF antibodies). (B) Genomic distribution of PLZF obtained from PLZF-Flag transduced 416b cell line (TSS: transcription start site defined from -1kb to +100bp; TES: transcription ending site defined from – 100bp to +1kb; UTR: untranslated region. (C) Gene Ontology (GO) analysis using GREAT tools of PLZF-peaks nearby genes (CC: cell cycle; DD: DNA damage; EP: epigenetics; CM: chemical modifications; IM: immunology). Gray scale values illustrate gene enrichment (i.e. number of genes observed/expected) for each GO term. (D) Venn diagrams showing the overlap between PLZF and histone marks.***P < 0.001 (hypergeometric test); NS not significant. (E) Scatter plots showing histone mark profiles in PLZF-Flag versus Empty vector conditions. (F) Venn diagrams showing the percentage of H3K27ac (upper) and H3K4me3 (lower) peaks modulated and/or bound by PLZF. (G) Integrative Genomic Viewer (IGV) screenshots of PLZF and H3K27ac in PLZF-Flag (PLZF) and Empty vector conditions (Empty).

Next, we analyzed the effect of PLZF over-expression on chromatin state. We performed differential analyses on histone signal in 416b in the presence (PLZF-Flag) or absence (Empty vector) of PLZF. Upon PLZF expression, H3K27ac and H3K4me3 ChIP-seq signals decrease, while H3K27me3 and H3K4me1 signals slightly increase (**Figure 1E and supplemental Figure S1B**) suggesting that the main chromatin effect of PLZF is to restrain already active genes through changes to the landscape of histone epigenetic marks. We found that there are twice as many downregulated H3K27ac sites than downregulated H3K4me3 bound by PLZF (**Figure 1F**), suggesting that PLZF preferentially targets H3K27ac. Decrease in H3K27ac level at PLZF-targeted genes was confirmed by Integrative Genomics Viewer (IGV) vizualisation (**Figure 1G**).

### Ectopic expression of PLZF modulates H3K27ac at enhancer regions in myeloid cells

H3K27ac is an activating mark found at promoters and marks active enhancer regions (33). Thus, we investigated whether the decrease in H3K27ac induced by PLZF-Flag was preferentially observed at one of these genomic regions. Since enhancer activity is cell-type specific and context sensitive, we determined enhancer regulatory regions using our 416b cell-specific ChIP-seq data. Promoter regions were defined as previously performed on human cells (27) but using the murine (mm10) genome. Enhancer regions were defined by the presence of H3K4me1 and absence of H3K4me3 and their activity was defined depending on the presence of additional histone marks (H3K27ac and H3K27me3). For example, active enhancers were H3K4me1+, H3K4me3- and H3K27ac+ (**Supplemental Figure S2A**). We observed that PLZF-induced H3K27ac decrease was mainly observed at enhancer regions with a higher significance and amplitude at active enhancers (**Figure 2A, Supplemental Figure S2B**), suggesting that PLZF would modify active enhancer activity. To further investigate whether PLZF is directly regulating enhancer activity, we analyzed PLZF genomic localization according to active enhancers. We observed that PLZF binds 40% of the active enhancers in PLZF-Flag 416b (**Figure 2B**) and that these enhancers were diminished in their H3K27ac level upon PLZF expression (**Figure 2C**). H3K27ac variations at PLZF bound active enhancers were validated by spike-in normalized ChIP-qPCR (**Figure 2D**). Next, we looked for the proportion of active enhancer regions that exhibit a decrease in H3K27ac (K27ac Down Active) upon PLZF expression and are bound by PLZF (PBDA, PLZF Bound Down Active, Table 2); we showed that more than 38% of K27ac Down Active enhancers were directly bound by PLZF (**Figure 2E**). GO analysis using the g:Profiler database confirmed that PBDA were enriched for genes associated with hematopoiesis (**Figure 2F**). Altogether, these results suggest that PLZF directly limits H3K27ac at enhancer regions controlling genes related to haematopoiesis.

**Figure 2:**
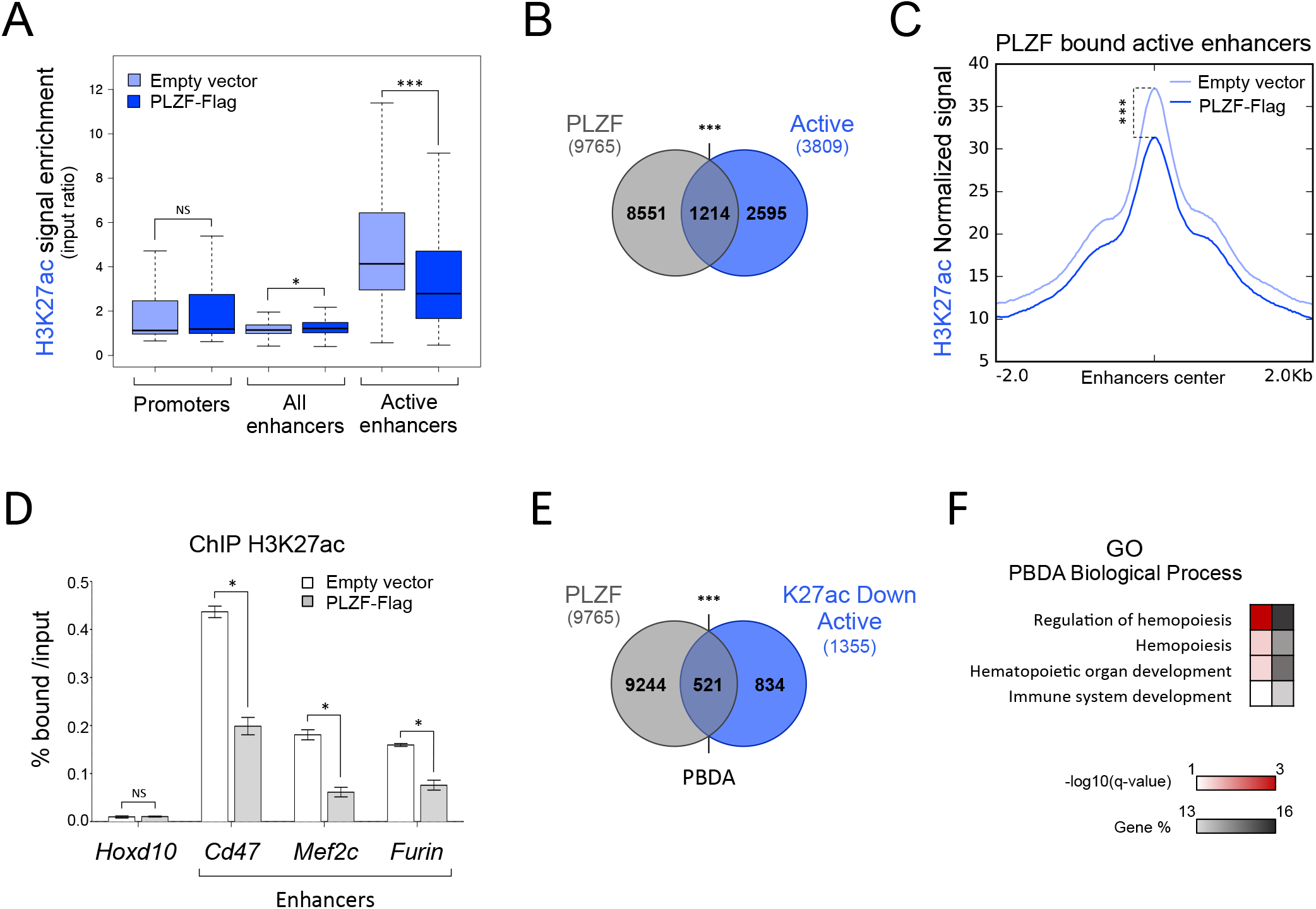
Ectopic expression of PLZF modulates H3K27ac at enhancer regions in myeloid cells. (A) Box plots showing H3K27ac enrichment (% H3K27ac bound/Input) in Empty vector and PLZF-Flag conditions at promoter, enhancer and active enhancer regions. *** *P* < 0.001 (Welch’s t.test); NS not significant. (B) Venn diagrams showing the overlap between PLZF peaks and active enhancers (Active) in PLZF-Flag condition. (C) Density plot profiles of H3K27ac normalized signal at PLZF-bound active enhancers in PLZF-Flag and Empty vector conditions. ***P < 0.001 (paired Welch’s t.test). (D) Spike-in ChIP-qPCR validation of H3K27ac variation at *Cd47, Furin* and *Mef2c* enhancers. *Hoxd10* is used as a negative control. Percentage of bound DNA over input are shown as a mean ±SD of two independent experiments (n=3, for each experiment). \**P* < 0.05 (Mann-Whitney test), NS not significant. (E) Venn diagrams showing the overlap between PLZF peaks and active enhancers with decreased H3K27ac level (K27ac Down Active) in PLZF-Flag condition. PBDA: PLZF-Bound Down Active. In figure B and E, \*\*\**P* < 0.001 (hypergeometric test); NS not significant. (F) Gene Ontology (GO) enrichment analysis on PBDA (intersect Figure E) using g:Profiler. Red scale indicates the p-value (-log10) and grey scale represents gene % ***(i.e.*** % of genes observed /total number of genes within each GO term).

### PLZF binding restrains enhancer activity

In order to evaluate the consequence of PLZF chromatin binding at enhancer regions, we measured chromatin accessibility by ATAC-seq and gene expression levels by RNA-seq in the absence and upon PLZF expression. First, ATAC-seq experiments showed that PLZF globally reduces chromatin accessibility (**Supplemental Figures S3A&B**). By analyzing PLZF-bound enhancer regions we revealed that when PLZF was expressed, these enhancer regions were reduced in their chromatin accessibility (**Figure 3A**). Decrease in ATAC-seq signal at some PLZF-bound enhancers was confirmed by IGV vizualisation (**Figure 3B**). These results suggest that PLZF binding at enhancers decreases their accessibility. Since enhancers can target genes located in their vicinity (34), we analyzed by RNA-seq gene expression according to the presence of PLZF in the associated enhancers. PBDA associated genes were separated into two groups depending on their PLZF ChIP-seq signal (low or high) and expression status of the two groups were compared to all genes. Results show that strong PLZF binding at enhancers (PLZF high) was associated with a significant lower gene expression in comparison to global expression (**Figure 3C**). When we compared RNA-seq counts from 416b cells with and without PLZF, we found that genes associated with PBDA consistently displayed reduced expression upon PLZF expression (**Figure 3D**), highlighting a direct effect of PLZF at enhancer region that restricts gene expression.

**Figure 3:**
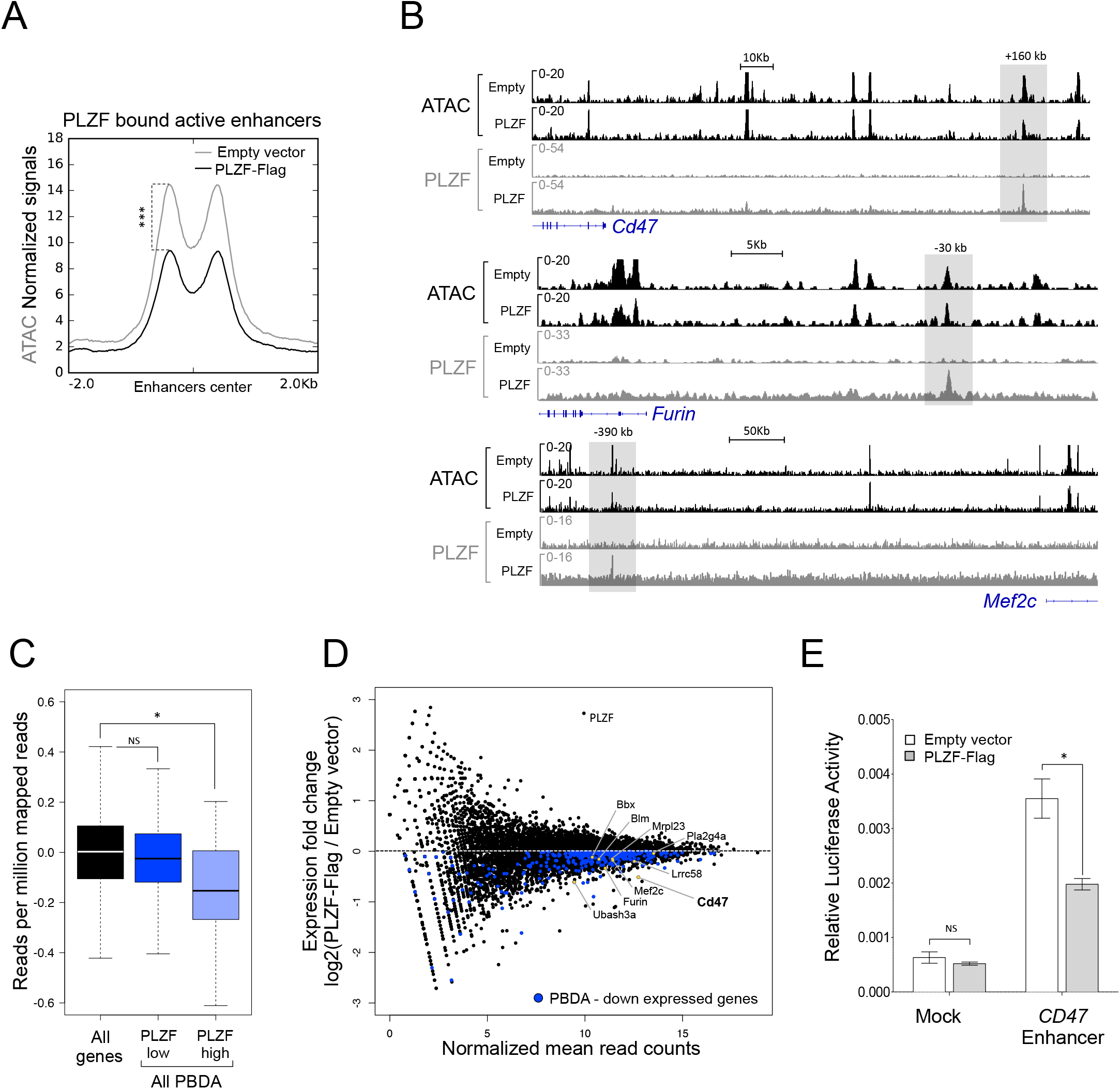
PLZF binding restrains enhancer activity. (A) Density plot profiles illustrating ATAC-seq normalized signal on PLZF-bound active enhancers in absence (Empty vector) or presence of PLZF (PLZF-Flag). \*\*\**P* < 0.001 (paired Welch’s t.test). (B) Representative Integrative Genomics Viewer (IGV) tracks of ATAC-seq, PLZF, signals at *Cd47, Furin* and *Mef2c* enhancers in PLZF-Flag versus Empty vector conditions. The grey box underlines the enhancers. (C) Box plots showing gene expression in PLZF-Flag condition of all genes and genes associated with PBDA with low (PLZF low) or strong PLZF (PLZF high) signals. PBDA: PLZF-Bound Down Active. \**P* < 0.05 (Welch’s t.test), NS not significant. (D) MA plot showing gene expression (RNA-seq) fold changes in PLZF-Flag *versus* Empty vector conditions (black dots). Blue dots represent downregulated genes associated with PBDA. Yellow dots underline some genes of interest. (E) Luciferase assay performed in 416b cells in absence (Empty vector) or presence of PLZF (PLZF-Flag) with control-LUC (Mock) or with CD47-enhancer-LUC *(CD47* Enhancer) reporters. Luciferase activity (FIREFLY: FF) was measured 8 h after transfection. FF values are normalized to RENILLA and expressed as a mean ±SD of three independent experiments. \**P* < 0.05 (Mann-Whitney test), NS not significant.

Finally, to recapitulate PLZF enhancer activity, we focused on one enhancer region bound by PLZF that regulates the CD47 gene (see **Figure 3B**). CD47 is involved in inflammatory response (35) and is recognized as an immune checkpoint for tumor evasion (36). We performed a Luciferase reporter assay using the previously described CD47 human enhancer (28) that we revealed to be bound by PLZF (extracted from Koubi et al., data). Luciferase activity monitoring showed that PLZF repressed the reporter activity under control of the CD47 enhancer (**Figure 3E**).

Altogether these data suggest that PLZF acts directly on enhancer regions to decrease their accessibility and activity.

### PLZF mutation increases H3K27ac at enhancers targeting genes related to metabolism

We previously showed that PLZF inactivation increases the Granulocytic Monocytic Progenitor (GMP) compartment after regenerative stress in the *Zbtb16^lu/lu^* mouse model (21). To question whether H3K27ac regulation by PLZF is linked to the GMP phenotype, we compared the chromatin landscape of *Zbtb16^lu/lu^* to WT GMPs after BM transplantation (**Figure 4A**). We profiled H3K27ac, H3K4me3, H3K4me1 and H3K27me3 distribution in WT and *Zbtb16^lu/lu^* GMPs. In accordance to our gain-of-function PLZF-Flag 416b cell line model, we showed that PLZF inactivation *in* vivo increased H3K27ac at the genomic level while H3K27me3 slightly decreased and the other epigenetic marks tested were modestly affected (**Figure 4B**).

**Figure 4:**
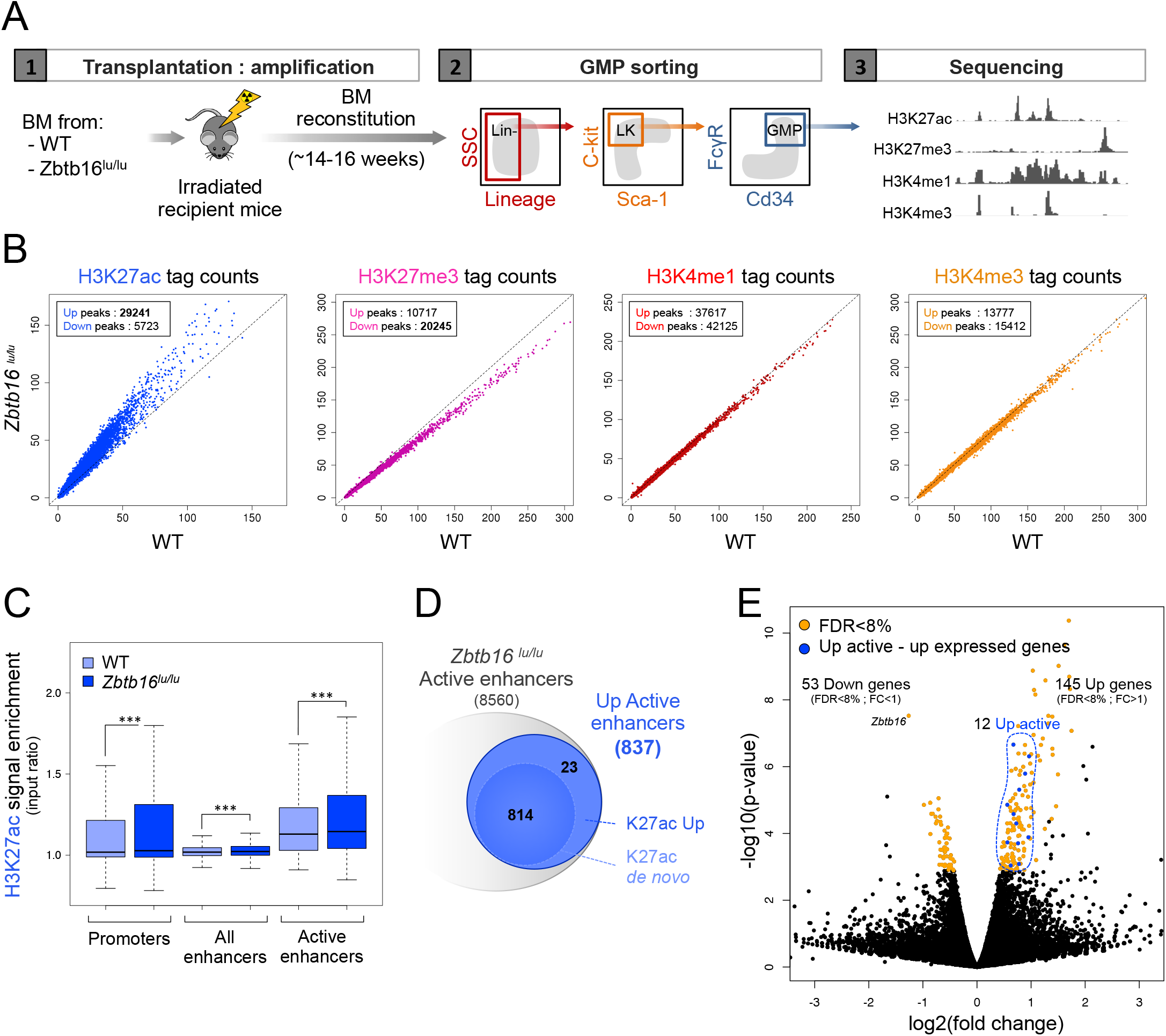
PLZF-mutation induced H3K27ac at enhancer regions in GMPs. (A) Experimental design used for generating histone mark ChIP-seq in wild type (WT) and PLZF mutant *(Zbtb16^lu/lu^)* Granulocyte-Monocyte Progenitors (GMPs). Lin-: Lineage negative, LK: c-kit^+^ and Sca-1^-^. (B) Scatter plots showing H3K27ac, H3K27me3, H3K4me1 and H3K4me3 signal in *Zbtb16^lu/lu^ versus* WT GMP. (C) Box plots showing H3K27ac enrichment (% H3K27ac bound/input) in WT and *Zbtb16^lu/lu^* GMPs at promoter, enhancer and active enhancer regions. \*\*\**P* < 0.001 (Welch’s t.test). (D) Venn diagrams showing the numbers of active enhancers in the *Zbtb16^lu/lu^* GMPs. K27ac Up: upregulated compared to the WT condition, K27ac *de novo*: not detected in the WT condition. (E) Volcano plot representing differential gene expression (RNA-seq) in *Zbtb16^lu/lu^* compared to WT. Yellow dot represents genes significantly modulated (false discovery rate (FDR) < 8%, fold change (FC) > 1 or < 1). Up Active enhancers associated with upregulated genes are highlighted in blue.

H3K27ac increase upon PLZF-mutation occurred both at promoter and enhancer regions (**Figure 4C**). In accordance with our overexpression model, we observe a stronger H3K27ac variation in mutant compared to WT when only active enhancers were considered (**Figure 4C, Supplemental Figure S4A**). In addition, a significant overlap was found between enhancers modified upon PLZF overexpression (Down Active) and upon PLZF mutation (Up Active) emphasizing the fact that PLZF globally restricts enhancer activity (**Supplemental Figure S4B**). Next, we investigated whether PLZF mutation favoured new H3K27ac enriched putative enhancers (“de *novo* active enhancers”) or was triggering already active enhancers. We showed that PLZF mutation increased H3K27ac at enhancer regions in both situation, when H3K27ac signal was detected in WT GMP (K27Ac Up: “already” active enhancers) but also when H3K27ac was not detected by our peak calling (K27Ac *de novo)* (**Figure 4D, Supplemental Figures S4C&D**). When considering all active enhancers identified in Mutant and WT GMPs (background, see materials and methods), *“de novo”* and “already” active enhancers modulated upon PLZF mutation exhibited a specific gene signature compared to unmodulated enhancers highlighting the specificity of PLZF on enhancer regions (**Supplemental Figure S4E, Table 3**). Indeed modulated enhancers were enriched for genes involved in response to stimulus and phosphorylation processes (**Supplemental Figure S4E**) whereas no significant gene signature was found for unaffected enhancers. To analyse whether changes in the epigenetic landscape would affect gene expression, we performed RNA-seq and compared the transcriptome of *Zbtb16^lu/lu^* to WT GMPs after BM transplantation. We showed that PLZF inactivation slightly affected gene expression (53 downregulated and 145 upregulated). Of these, 10% of significantly upregulated genes associated with Up Active enhancers exhibited higher expression upon PLZF inactivation (**Figure 4E**). Altogether, these results show that PLZF inactivation results in an increase in H3K27ac at enhancer regions.

### Aging of GMPs is marked by an accumulation of H3K27ac at enhancer regions

We previously showed that PLZF limits some of the HSC aging features (21). Thus, we asked whether the PLZF specificity on enhancers was linked to hematopoietic aging. For this purpose, we investigated H3K27ac modulation in GMP compartment of aged mice. Comparison of H3K27ac level in Old and Young WT GMPs revealed a global H3K27ac increase in Old GMPs (**Figure 5A**). Comparable to what we observed in *Zbtb16^lu/u^* GMPs, this increase was significant at active enhancer regions (**Figure 5B**). Among the 2307 putative active enhancers found in old GMPs, 816 gained H3K27ac upon aging. This gain of H3K27ac was mostly observed in *“de novo”* active enhancers (**Figure 5C**). Interestingly, among the 1283 genes associated with these enhancers that gain H3K27ac upon aging (Old WT Up Active), 27% (346 genes, “common”) were also affected by PLZF deletion *(Zbtb16^lu/lu^* Up Active) (**Figure 5D, Table 4**). Interestingly, 86% of these conjointly modulated enhancers were potentially bound by PLZF (**Supplemental Figure S5A**). When considering all active enhancers identified in Young, Old, WT and Mutant GMPs as background enhancer set (see materials and methods), these conjointly regulated enhancer regions (common) were enriched in genes associated with metabolism (**Figure 5E**). They were also related to abnormal inflammatory response when extending the analysis to associated-human phenotype (**Figure 5E**). Interestingly, considering the same background enhancer set as above no significant biological processes were found for enhancers modulated exclusively by aging (937 genes) or by PLZF deletion (1068 genes). As enhancers modulated upon aging appeared to be under the control of PLZF, we monitored by qPCR the expression of PLZF in Young and Old GMPs. We showed that Young and Old GMPs exhibited similar ***Plzf*** expression levels (**Supplemental Figure S5B**), suggesting that aging may alter PLZF chromatin activity rather than its expression level.

**Figure 5:**
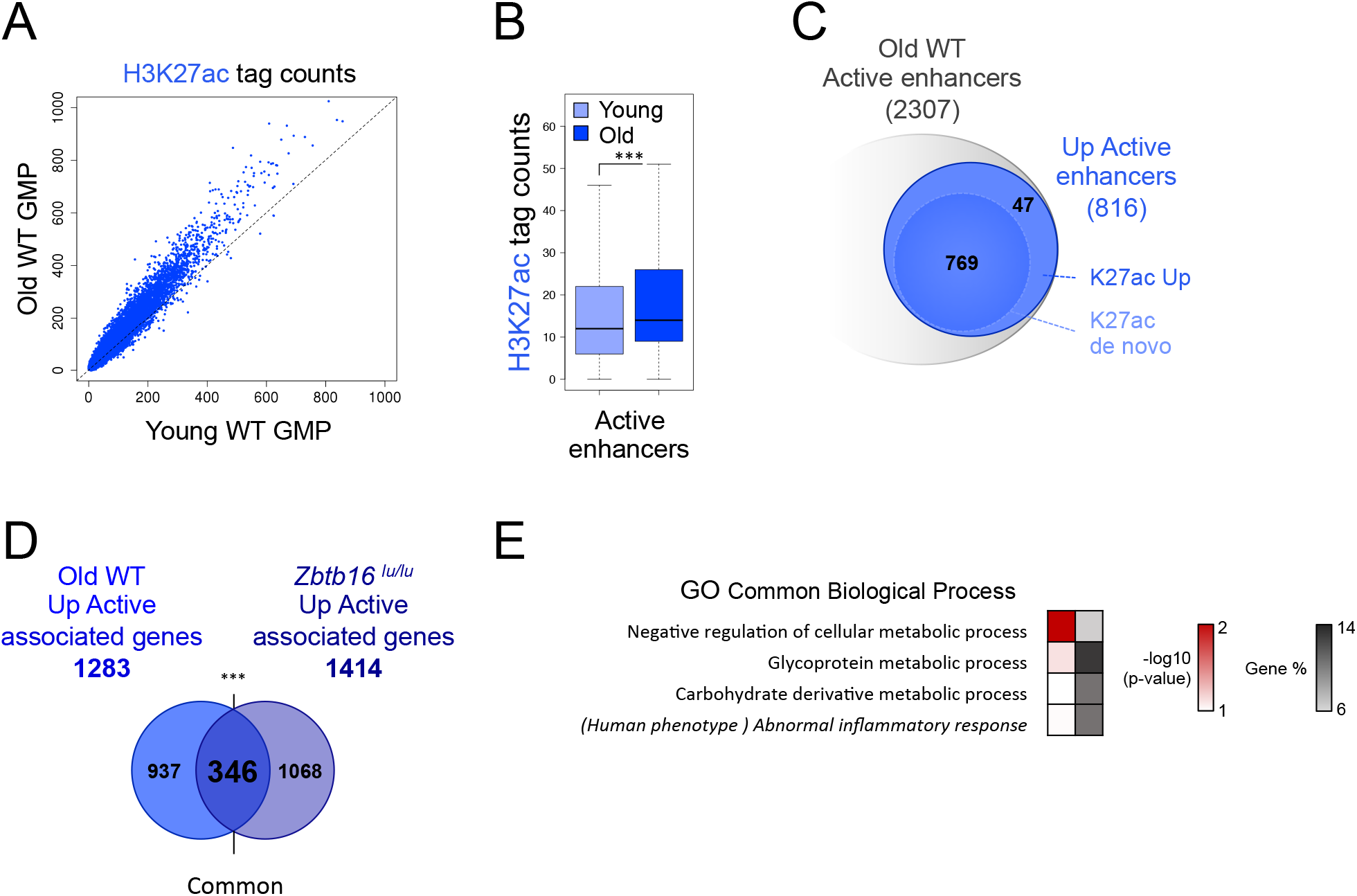
Aging of GMPs is marked by an accumulation of H3K27ac at enhancer regions. (A) Scatter plots showing H3K27ac profiles in Old *versus* Young GMPs. (B) Box plot showing H3K27ac level at active enhancers in Old *versus* Young GMPs. \*\*\**P* < 0.001 (Welch’s t.test). (C) Venn diagrams showing the numbers of active enhancers in Old WT GMPs. K27ac Up: upregulated compared to the Young condition, K27ac *de novo*: not detected in the Young condition. (D) Venn diagram showing the overlap (common) between genes associated with active enhancers upregulated during aging (Old Up Active associated genes) and genes associated with active enhancers upregulated in *Zbtb16^lu/lu^* GMPs (Zbtb16^lu/lu^ Up Active associated genes) \*\*\**P* < 0.001 (hypergeometric test). (E) Gene Ontology (GO) analysis and associated human phenotype of “common” enhancers that change in aged and *Zbtb16^lu/lu^* conditions (intersect Figure D) using g:Profiler. Red scale indicates the p-value (-log10) and grey scale represents gene % (*i.e.* % of genes observed /total number of genes within each GO/KEGG term).

Altogether, these results show that increase in H3K27ac level at enhancer regions controlling metabolism-related genes may be a hallmark of hematopoietic progenitor aging that is controlled by PLZF activity.

## DISCUSSION

PLZF is a transcription factor involved in multiple facets of cell biology. Here we reveal a new molecular function of PLZF in limiting H3K27ac at enhancer regions that may restrict their activity. H3K27ac is a well-defined marker of active enhancers (33), and it is essential for their activation (37). Thus, by showing that PLZF controls H3K27ac level in murine progenitor cells, we highlight a novel PLZF chromatin activity that is merely to restrict active enhancers. Gain of function experiments support a direct effect of PLZF on H3K27ac and enhancer activities. This goes in line with previous studies showing that PLZF recruits HDAC complexes at targeted promoters and locally reduced histone acetylation (25,26,38). However, the global increase of H3K27ac, not restricted at PLZF binding sites, that is observed in PLZF-mutated myeloid progenitors also suggests that PLZF has an effect on acetyl regulation beyond its direct chromatin activity. This close relation between PLZF and acetylation is not only supported by our ChIP-seq experiments that revealed enrichment of PLZF at genes involved in protein acetylation but also by the fact that PLZF transcriptional activity is itself modified by acetylation (39, 40).

Due to its affinity with Polycomb group proteins (23, 24), PLZF was first described as a transcriptional repressor. The repressive chromatin activity of PLZF has been challenged previously by genomic data analyses showing the presence of PLZF on regulatory elements of active genes (12,27,41). Here again, analyzing murine myeloid progenitors, we found that PLZF is mainly present at already active chromatin. However, by modulating its expression in myeloid progenitors (overexpression in 416b cells and mutation in GMPs), we demonstrate that the presence of PLZF reduces the H3K27ac active mark and that this reduction was marked at active enhancers emphasizing its repressive activity. This suggests that PLZF limits enhancer activity through acting as a “brake” on these regulatory elements. This notion of “brake” to ensure appropriate enhancer activity was previously proposed for RACK7/KDM5C complex that controls enhancer over-activation by modulating histone methylation (42).

This “brake” function on enhancer activity fits well with PLZF hematopoietic function: PLZF has been described to modify chromatin to restrain the production of pro-inflammatory cytokines in case of a bacterial infection in bone marrow-derived macrophages (38). Interestingly, this restriction involves acetylation of PLZF itself that promotes the assembly of a repressor complex incorporating HDAC3 and the NF-kB p50 subunit that limits the NF-kB signalling (40).

It is also striking to notice that the strong gain of H3K27ac at enhancer regions in PLZF-mutant GMPs has little repercussion at the transcriptional level. Differential gene expression analyses revealed that only 12 H3K27ac-UP associated genes are more expressed in the PLZF mutant condition. This apparent discrepancy between chromatin structure and gene expression is not without precedent (43). Chromatin modifications may precede, or prime chromatin for, transcriptional changes. It is also reported that chromatin structure identifies hematopoietic cell population better than gene expression profile (44). This suggests that the absence of PLZF would prime chromatin to be able to respond to cell stimuli.

In line with this idea, we highlighted a clear increase in H3K27ac, marked at enhancer regions in aged myeloid progenitors, comparable to what we observed in PLZF-mutant progenitors. There is a growing body of evidence demonstrating that epigenetic changes are central to age-associated tissue decline (45). The main common aging-chromatin feature is a global loss of closed-chromatin, characterized by a decrease in H3K9me3 (46, 47), a gain in H3K4me3 specifically at promoters of self-renewing genes and a global DNA hypomethylation (48). While histone acetylation, *via* the study of two deacetylases Sirt1 and Sirt3, was previously shown to play a role under stress conditions in aging mouse HSC (49, 50), the direct assessment of H3K27ac in hematopoietic cell aging has not before been addressed. Here, we found that H3K27ac is a hallmark of aged GMPs and observe a loose chromatin structure at genes involved in metabolic processes while aging. Indeed, metabolic deregulation was reported in several aging models (21, 51). Interestingly, aged-related dysfunction of metabolism could in turn affect epigenetics such as deregulation of NAD+ production upon aging that leads to enzymatic deficiency of the NAD-dependent histone deacetylase Sirt1 (52).

PLZF expression does not decline during aging but its activity may change. Recent studies have highlighted that epigenetic cofactors having post-translational modification properties could also modify PLZF itself and influence its transcriptional competence. PLZF transcriptional repression required the acetyltransferase activity of P300 (53). More recently, PLZF was shown to interact with the histone methyltransferase EZH2, which could methylate PLZF affecting its stability and its transcriptional activity (27, 54). In addition, HDAC7, a key factor of the innate effector programming of iNKT cells, changes PLZF activity underlying the tight connection between specific transcription factor activity and post-translational modification of the epigenetic machinery (55).

In conclusion, we have elucidated a novel function for PLZF, a known epigenetic regulator, in restricting enhancer acetylation of genes involved in hematopoietic aging.

## ACCESSION NUMBERS

ChIP-seq data (FastQ and Bigwig files) are in the process of being deposited in the NCBI Gene Expression Omnibus (GEO; http://www.ncbi.nih.gov/geo/).

## SUPPLEMENTARY DATA

### FUNDING

This work was supported by Institut Thématique Multi-Organisme-cancer (grant number C13104L to ED and CG; P036560 to ED and LN), by l’Institut National du Cancer (grant number 20141PLBIO06-1 to ED, CG and JV), by the Fondation de France to MP, by the JSPS program to MP and by Aix Marseille University to LH.

## Supporting information

Supplemental informations

## ACKNOWLEDGMENTS

Authors thank N. Carbuccia, J. Adelaide and A. Saraya for library preparation and sequencing of ChIPed DNA and Dr Takayama for his help for ATAC-sequencing. Authors gratefully acknowledge P.A. Betancur for giving them *CD47* enhancer constructs and Dr. Göttgens for giving them 416b cell line. Authors gratefully acknowledge cytometry, animal and DISC platforms of CRCM.

