## Supplemental informations for "PLZF limits enhancer activity during hematopoietic progenitor aging"

### Figure S1

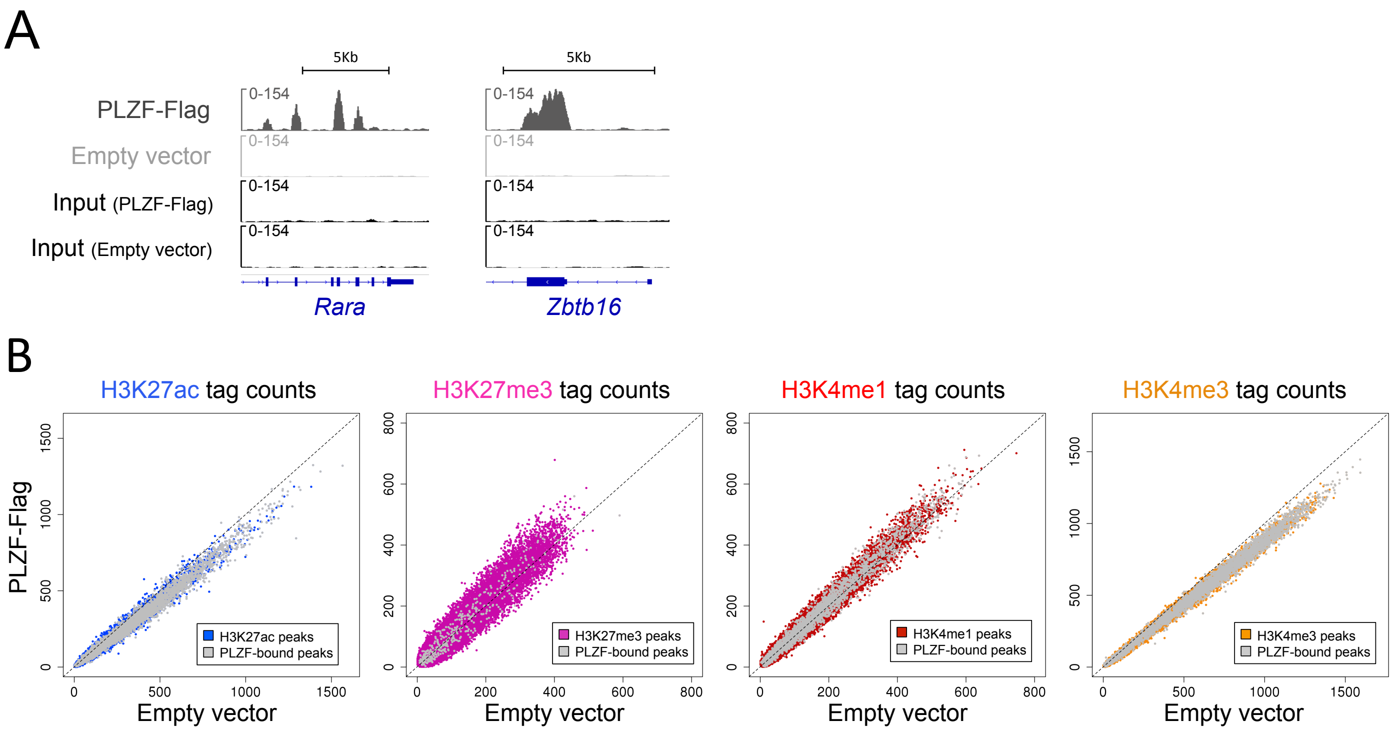


**Figure S1. Related to figure 1**

(A) Representative Integrative Genomics Viewer (IGV) screenshot of PLZF peaks (PLZF-Flag) on previously described targeted genes. (B) Scatter plots showing histone mark profiles in PLZF-Flag *versus* Empty vector conditions. Grey dotes represent sites bound by PLZF.

### Figure S2

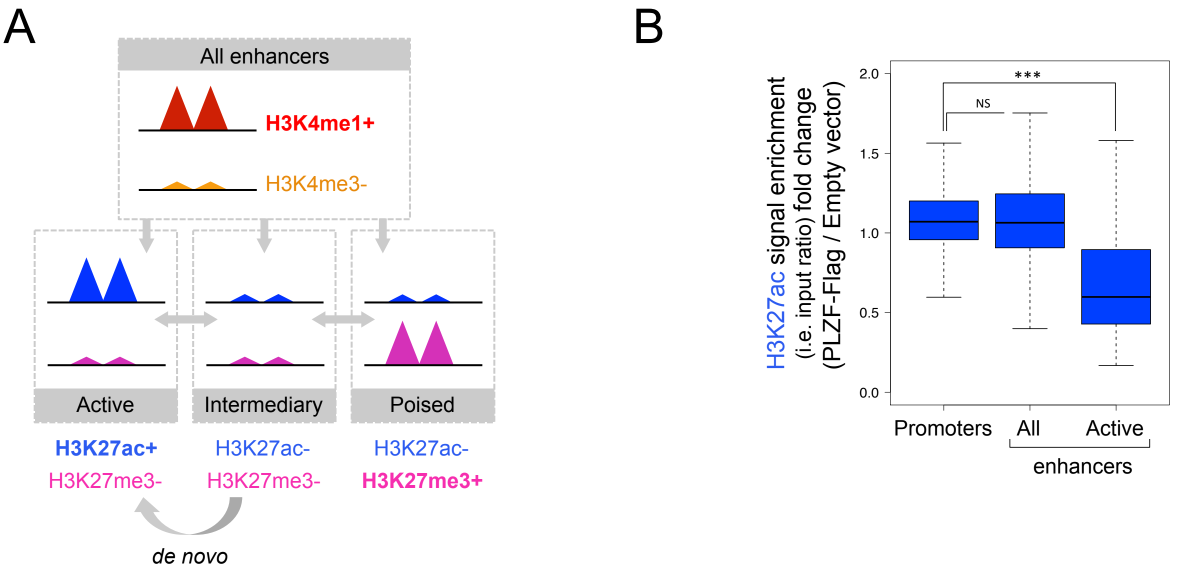


**Figure S2. Related to figure 2**
(A) Schematic representation of enhancers. Enhancer regions are determined with H3K4me1 (high) and H3K4me3 (low) levels. Intermediary enhancers lack H3K27ac and H3K27me3 but could gain these histone marks. Active enhancers are H3K4me1+, H3K4me3-, H3K27ac+), poised enhancers are H3K4me1+, H3K4me3-, H3K27me3+. Enhancers gaining H3K27ac are defined as "*de novo* active enhancers”. (B) Box plot showing H3K27ac enrichment (% H3K27ac bound / Input) fold change (PLZF-Flag / Empty vector) at promoter, enhancer and active enhancer regions. . ***$P < 0.001$ (Welch’s t.test); NS not significant

### Figure S3

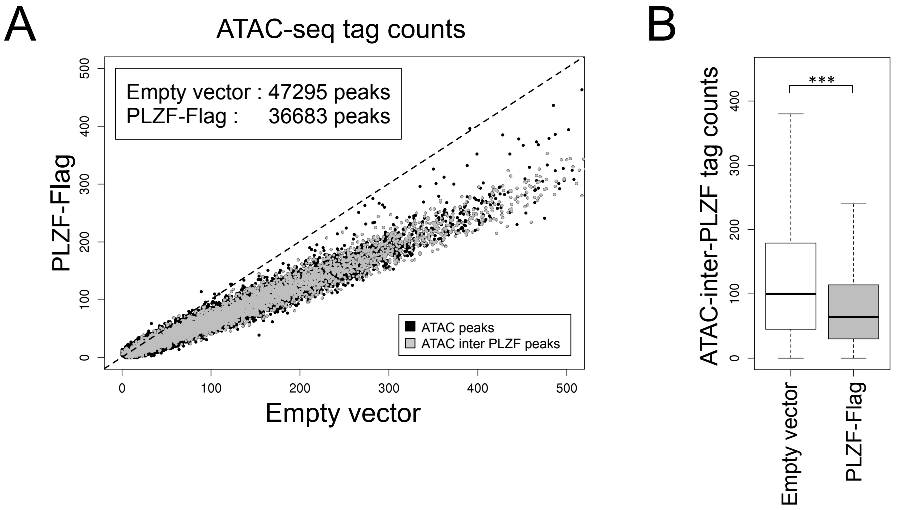


**Figure S3. Related to figure 3**

(A) Scatter plots showing ATAC-seq signal in PLZF-Flag *versus* Empty vector conditions. Grey circles represent sites bound by PLZF. (B) Box plots showing ATAC-seq signal at PLZF-bound sites in Empty vector and PLZF-Flag conditions. ***$P$ < 0.001 (Welch's t.test).

### Figure S4

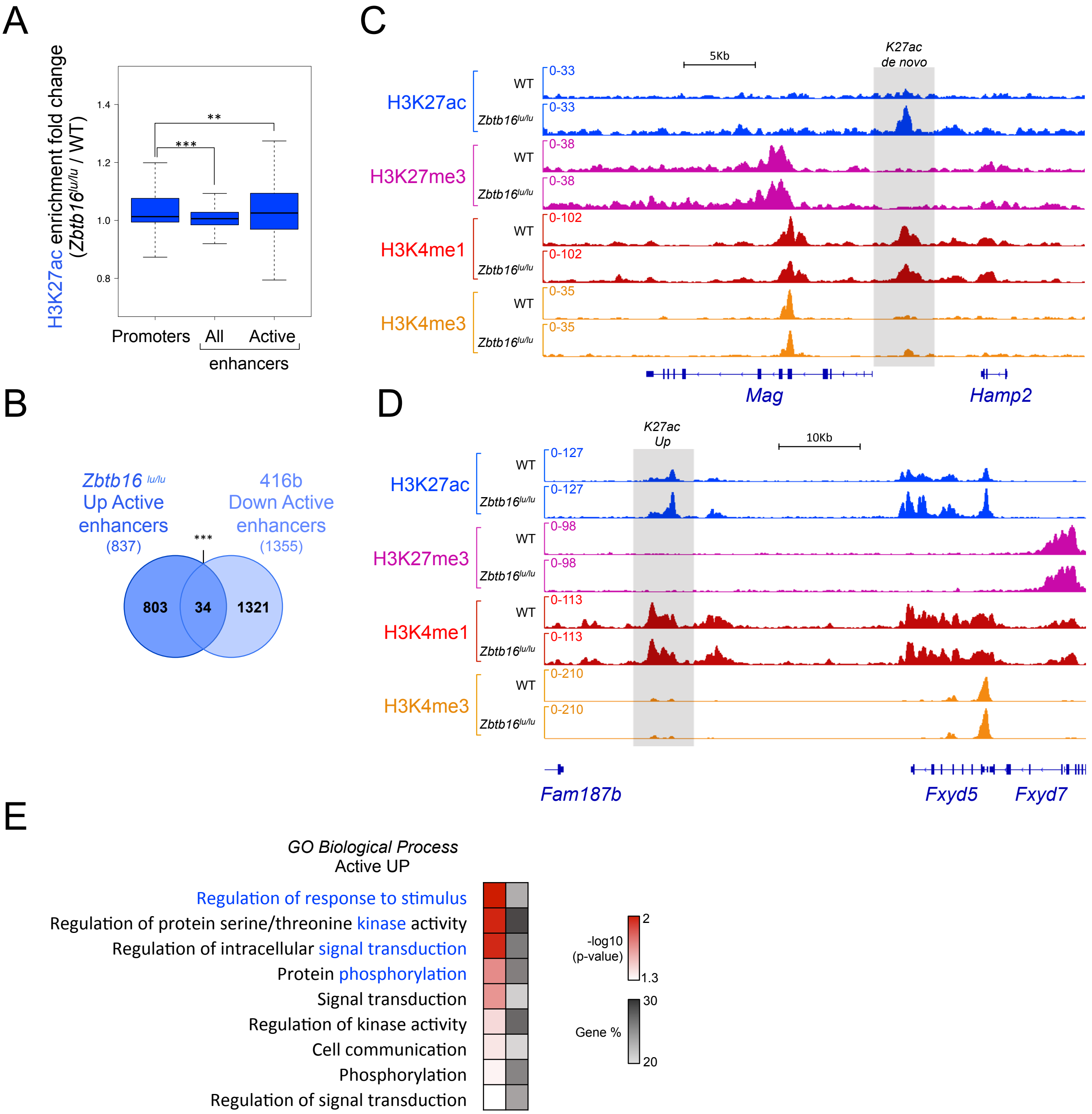


**Figure S4. Related to figure 4**
(A) Box plot showing H3K27Ac enrichment (% H3K27ac bound / Input) fold change (*Zbtb16*$^{lu/lu}$ / WT) at promoter, enhancer and active enhancer regions. ***P* < 0.01; ****P* < 0.001 (Welch's t.test). (B) Venn diagrams showing the intersection between enhancers modified upon PLZF over expression (Down Active, from 416b) and upon PLZF mutation (Up Active, from GMPs). ****P* < 0.001 (hypergeometric test) (C-D) Representative Integrative Genomics Viewer (IGV) tracks of H3K27ac, H3K27me3, H3K4me1 and H3K4me3 in WT and *Zbtb16*$^{lu/lu}$ GMPs at enhancer regions. The grey box underlines the region of interest. (E) Gene Ontology (GO) enrichment of genes associated with enhancers that increase (Active UP: K27ac *de novo* and K27ac Up). Red scale indicates the p-value (-log10) and grey scale represents gene % (*i.e.* % of genes observed /total number of genes within each GO term).

### Figure S5

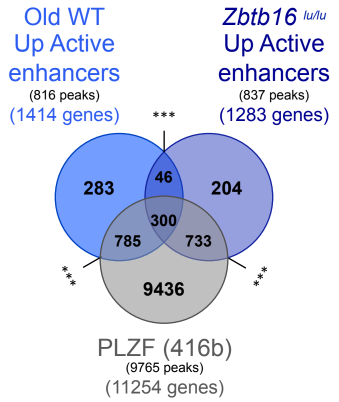


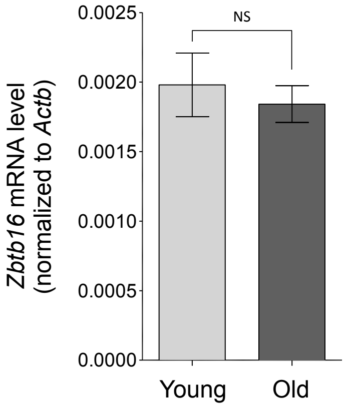


**Figure S5. Related to figure 5**

(A) Venn diagrams showing the intersection between enhancers modified upon aging (Old WT Up Active), enhancers modified upon PLZF mutation (*Zbtb16*$^{lu/lu}$ Up Active), and PLZF peaks from 416b cells. ***$P$ < 0.001 (hypergeometric test). (B) RT-qPCR showing *Zbtb16* expression levels in Young and Old GMPs. mRNA values are normalized to *Actb* and expressed as a mean ±SD (n=3); NS not significant.
